# Acute nicotine exposure produces a spatially organized transcriptional response in mouse olfactory cortex

**DOI:** 10.64898/2026.11.04.764395

**Authors:** Mina Griffioen, Andrew Lombardi, Ryan Milstead, Curtis Borski, Shiying Zou, Jerry Stitzel, Marissa Ehringer, Charles Hoeffer

## Abstract

Nicotine acts on nicotinic receptors throughout the forebrain. Most molecular studies of its effects, however, have focused on reward circuitry and have used bulk tissue, which cannot show which cells respond or how responding cells are arranged relative to one another. We previously found that a single nicotine exposure activates hippocampal astrocytes through the astrocyte-expressed kinase AKT2. Here we asked which cells respond to nicotine in the anterior olfactory cortex, a cholinergic-modulated region that encodes odor associations, and whether their astrocytic surroundings change. Male mice received a single intraperitoneal injection of nicotine or saline, and coronal sections spanning the anterior olfactory nucleus, piriform cortex and adjacent isocortex were sequencing 24 h later using Visium HD. Differential expression on nucleus-based segmentation was concentrated in olfactory excitatory neurons. Pathways centered on calcium-handling, synapse-assembly and extracellular-matrix programs, together with a lasting reduction of the immediate-early gene *Egr1*. Within the piriform region, the response localized to a continuous population of superficial pyramidal neurons expressing *Cartpt* and *Penk*. These neurons upregulated synapse-assembly, receptor-regulation and proteoglycan programs without changes to cell-type proportions. Nicotine also reduced expression of *Sparcl1*, which encodes astrocyte-secreted synaptogenic protein hevin, within one cell diameter of these neurons. Our study shows that a single nicotine exposure leaves a spatially organized transcriptional program on olfactory circuitry, centered on one neuronal population and the tissue at its border. These results identify candidate cells, genes and tissue locations for testing how nicotine alters sensory cue circuits.

## Introduction

Tobacco use remains a leading preventable cause of death worldwide, killing more than 8 million people each year (1). Nicotine is the principal psychoactive component of tobacco, and most molecular studies of its actions have focused on mesolimbic reward circuitry. Nicotine also acts throughout the forebrain through widely distributed nicotinic acetylcholine receptors, and acute nicotine enhances attention and memory in humans (2,3). Approved pharmacotherapies (nicotine replacement, bupropion, and varenicline) help, but most people who use them do not achieve long-term abstinence, and relapse is common (4), underscoring the need to understand how nicotine alters brain circuits beyond the reward pathway. The cell types through which a single nicotine exposure alters gene expression outside reward pathways, and how those changes are organized within tissue, remain poorly understood.

The piriform cortex and anterior olfactory nucleus (AON) receive dense cholinergic input from the basal forebrain, and cholinergic modulation of these regions supports olfactory learning and memory (5,6). Within piriform layer 2, semilunar cells receive strong direct input from the olfactory bulb, whereas superficial pyramidal cells integrate bulbar input with recurrent associational input and project to frontal cortex (7,8). Nicotine engages this system: a single injection induces *c-fos* mRNA in piriform cortex (9), chronic oral nicotine improves olfactory memory through TRPA1 (10), and repeated nicotine exposure alters glutamate-associated network connectivity involving piriform cortex (11). Piriform glutamatergic neurons also vary continuously rather than forming discrete types (12), making tissue position an informative addition for defining them. Nicotine also enhances glutamatergic release (13) and facilitates synaptic potentiation (14). Nicotine’s effects on olfaction also differ from those of smoking. Cigarette smoking impairs olfactory function in a dose-related manner(15,16), whereas nicotine itself has been reported to improve olfactory memory in mice (10), suggesting that nicotine can act on olfactory circuits independently of the other constituents of tobacco smoke. These studies support studying the olfactory cortex as a site where nicotine may engage cholinergic learning circuitry, and glutamatergic signaling.

Nicotine also acts directly on glia. Astrocytes express functional α7 and α4β2 nicotinic receptors, whose activation elevates astrocytic calcium (17), releases factors that recruit AMPA receptors to glutamatergic synapses (18), and remodels astrocyte processes (19,20). Our previous work showed that astrocytes are among the cells that respond to nicotine. We found that acute nicotine exposure increased astrocyte morphological complexity, that chronic nicotine had the opposite effect, and that the astrocyte-restricted kinase AKT2 modulated these responses and nicotine conditioned place preference (19). Bulk RNA sequencing of the hippocampus after acute nicotine in that study identified differentially expressed genes associated with nicotine-related smoking initiation genome wide association study (GWAS) loci. However, bulk sequencing averages across cell types and cannot determine spatial position, making it unable to identify which cell types carry a transcriptional response or whether neurons and the astrocytes surrounding them change together. Single-nucleus studies have begun to resolve cell-type-specific responses to nicotine in the ventral tegmental area (21), but dissociating tissue also discards position, and to our knowledge nicotine’s effects on the brain have not been examined with spatial transcriptomics.

Here we used Visium HD with nucleus-based cell segmentation to profile the anterior olfactory cortex and adjacent isocortex 24 h after a single intraperitoneal injection of nicotine or saline, the same paradigm as our previous study. We asked which cell types respond, where the responding cells are located, and whether the astrocytic environment around them changes. We found that the response was concentrated in olfactory excitatory neurons and was localized to a population of superficial pyramidal neurons expressing *Cartpt* and *Penk* and was accompanied by a local reduction of the astrocyte-secreted synaptogenic gene *Sparcl1* at their border.

## Methods

### Animals and nicotine exposure

Male wild-type (C57BL/6) mice 2-6 months of age were used to determine spatial gene expression signatures exclusive to nicotine exposure. Mice received a single intraperitoneal (i.p.) injection of nicotine (0.2mg/kg) (N = 2), or an equal volume of 0.9% saline vehicle (N = 2). Mice were euthanized 24 h later using cervical dislocation, and brains were rapidly extracted and flash-frozen in isopentane chilled on dry ice. Frozen brains were embedded in optimal cutting temperature (OCT) compound and stored at −80°C until sectioning. Animal care and treatment procedures were approved by the University of Colorado Boulder IACUC and follow NIH Guidelines, mice were kept in cages with sex-matched littermates on a 12:12 h light:dark cycle with an ambient temperature ranging from 22-24 °C, and food and water were given *ad libitum*.

### Tissue sectioning and Visium HD library preparation

Coronal sections (20 µm) were cut on a cryostat at −20 °C, mounted onto positively charged slides, and stained with hematoxylin and eosin (H&E), and imaged at 20X with EVOS M7000. Tissues were mounted onto 10x Genomics Visium HD CytAssist (10x Genomics, #1000669), targeting anterior sections containing the piriform/olfactory and adjacent isocortical areas. Visium HD libraries were prepared according to the manufacturer’s protocol (10x genomics #1000667). Libraries were sequenced to a depth of ∼500-600M read pairs per section.

### Read processing and cell segmentation

Raw sequencing reads were processed with Space Ranger (10x Genomics v4.0.1) against the mouse reference genome GRCm39. Rather than the default 2 µm binned output, near single-cell resolution was obtained using Space Ranger’s nucleus-based cell segmentation pipeline, which detects nuclei from the H&E image and expands segmentation boundaries to assign transcripts to individual cells (“polygons”). Segmented per-cell feature-barcode matrices, cell coordinates, and scale factors were used for all downstream analyses.

### Quality control, integration, and clustering

Segmented outputs from all four sections were loaded into Seurat (v5.1.1); R v4.4.1) as spatial polygon assays. Low-quality cells were removed using thresholds on transcript counts and detected genes (>100 detected genes, >200 transcripts, and <15% mitochondrial transcripts per cell, applied identically to all sections; retaining 450,785 of 477,673 total cells). To integrate the four sections while controlling for section-level technical variation, we used a sketch-based workflow: a representative subset of n = 60,000 cells were sampled across sections (n = 15,000 cells per section), log-normalized, and used for shared feature selection. Batch effects were corrected across sections with Harmony (22), and cells were clustered on the Harmony-corrected embedding using the Louvian algorithm (resolution 0.5), yielding 17 clusters. Cluster labels were then projected from the sketched cells onto the full dataset. Uniform Manifold Approximation and Projection (UMAP) was computed on the Harmony embedding for visualization.

### Cell type annotation

Cells were annotated with the Allen Brain Cell (ABC) atlas whole-brain taxonomy tool (hierarchical mapping to the 10x whole-mouse-brain taxonomy, CCN20230722; (23)), and cluster identities were assigned from the consensus of cell-level mapping together with canonical marker genes. Atlas mapping confirmed that the captured tissue comprised anterior isocortical, olfactory, and piriform/entorhinal cortical populations together with associated glial and vascular cell types. One low-count cluster (median 531 transcripts per cell, the lowest of any cluster and roughly one-third to one-half of the per-section medians of 1,125-1,710) expressing a mixture of neuronal and astrocyte-associated transcripts was determined to represent a segmentation artifact of neuronal-astrocyte boundary assignments rather than a bona fide cell population, and was excluded from differential and inferential analyses. A separate astrocyte-containing cluster was subclustered to resolve astrocytes from adjacent meningeal and mural cells. For plotting and comparison, cell-type labels were stored in metadata; non-biological or confounded populations (segmentation-artifact cluster, sample-driven populations, meningeal and mural cells) were excluded from condition comparisons.

### Piriform/olfactory region-of-interest definition

To focus on the piriform/anterior olfactory response, a region of interest (ROI) was manually drawn on each section to contain the piriform (PIR), entorhinal (ENTl), and anterior olfactory nucleus/taenia tecta (AON/TT) areas. Cells falling within the ROI were re-annotated at higher resolution into region-specific populations, including a *Cartpt/Penk*-expressing excitatory compartment (PIR-ENTl *Cartpt/Penk*), a PIR-ENTl/AON glutamatergic population, AON-TT populations, and local inhibitory, astrocytic, oligodendrocyte, OPC, and mural/vascular populations. ROI cells (158,478) were re-normalized, 3,000 variable genes were used for principal component analysis, sections were integrated with Harmony, and cells were clustered on 30 components (resolution 0.4, chosen after testing at 0.3 and 0.8, for separation of known piriform and AON marker genes).Three astrocyte clusters with nearly identical depth-normalized marker profiles, differing mainly in transcript depth, were merged into a single astrocyte population. Differences in cell-type proportions between conditions were tested with propeller (24).

### Differential expression (pseudobulk)

Animal (N = 2) was used as the unit of replication to avoid using spatial polygons within a section as independent biological replicates. For each cell population, raw transcript counts were aggregated across cells to a single pseudobulk profile per animal (Seurat AggregateExpression; (25)). Differential expression between nicotine and saline was tested with DESeq2 using a ∼condition design with saline as the reference, applying apeglm log-fold-change shrinkage for whole-tissue comparisons (Love et al., 2014; Zhu et al., 2019); ROI comparisons used unshrunken estimates. Genes were retained if they exceeded a minimum aggregate count (rowSums ≥ 10), and mitochondrial (*mt-*) and ribosomal (*Rps/Rpl*) genes were excluded from downstream summaries and visualizations. Genes with Benjamini-Hochberg adjusted p < 0.05 were considered differentially expressed. The same pseudobulk framework was applied to both whole-tissue cell types and the ROI-defined populations.

### Gene set enrichment analysis, spatially variable and differential spatial pattern genes

Gene set enrichment analysis (GSEA) was performed on the *Cartpt/Penk* and whole-tissue piriform differential expression analysis results using clusterProfiler (gseGO; Wu et al., 2021) against Gene Ontology Biological Process terms, with genes rank by the Wald statistic after excluding mitochondrial and ribosomal genes. Terms with adjusted p < 0.05 were considered significant. A targeted test of the KEGG nicotine addiction gene set (mouse orthologs of hsa05033) was performed with fgsea (26) on the same rankings.

Spatially variable genes (SVGs) were identified per section with SPARK-X (27) on segmented cell coordinates and normalized expression, restricting to genes detected in at least 1% of cells. To identify genes whose spatial variability was condition-dependent, and due to small sample size, a strict replicate-consistency criterion was applied. Genes were considered nicotine-gained if it was spatially variable (adjusted p < 0.05) in both nicotine sections and neither saline section, and vice versa. Differential spatial pattern (DSP) genes whose expression pattern across spatial domains differs between conditions were identified using the DESpace framework (28). Cell-type populations were treated as spatial domains and pseudobulk counts were aggregated per animal x domain and modeled in edgeR with a domain x condition interaction term (∼domain + condition + domain:condition, quasi-likelihood F-test on interaction coefficients). Genes with interaction of FDR < 0.05 were considered DSP genes.

### Cartpt/Penk spatial gradient analysis

To test whether gene expression varied with spatial position relative to the *Cartpt/Penk* compartment and whether nicotine altered that relationship, the physical distance from each PIR-ENTl/AON glutamatergic cell to the nearest *Cartpt/Penk* cell was computed and converted from full-resolution pixels to micrometers using section-specific scale factors. Cells were assigned to physical distance bins (0-12, 12-24, 24-48, 48-96, and 96-192 µm), where the first bin approximates one cell diameter, doubling bin widths keep cell numbers comparable across distances, and cells beyond 192 µm were excluded. The model uses each animal-bin’s median distance as a continuous covariate to avoid results depending on exact bin edges. The design adapts the spline-interaction approach used for time-course experiments in limma/edgeR (29) to a spatial covariate, methods that relate expression to continuous distance from a tissue feature (30). Raw counts were aggregated per animal x distance bin. A replicate-aware model was fit in edgeR (robust quasi-likelihood) of the form ∼sample + ns(log2(distance), df = 2) + condition:ns(log2(distance), df =2), where a natural cubic spline captured the expression-distance relationship. A joint quasi-likelihood F-test of the condition x spline interaction terms tested whether the shape of the expression-versus-distance curve differed between nicotine and saline (10,861 genes, those passing edgeR filterByExpr in the animal x distance-bin pseudobulk after removal of mitochondrial and ribosomal genes).

As orthogonal, within-section validation of spatial structure, boundary-proximity analyses were performed using SpNeigh (v1.1.2) (31). *Cartpt/Penk* cells defined the boundary, inverse-distance boundary weights were computed for PIR-ENTl/AON glutamatergic cells (computeBoundaryWeights), and spatial expression trends were tested with runSpatialDE (df = 2) on log1p CP10k-normalized expression after filtering to genes detected in ≥5% of cells and removing mitochondrial/ribosomal genes. Gaussian-process spatial variability within sections was additionally assessed with SPADE (v0.99.0)(32) on µm-scaled coordinates. These within-section analyses establish the existence of the spatial gradient and are distinct from the replicate-aware edgeR test of a nicotine effect on the gradient.

### Cell-cell communication analysis

Cell-cell communication was inferred with CellChat (v2.2.0.9001)(33) using the mouse ligand-receptor database (CellChatDB.mouse). Spatially-informed inference was performed per section using segmented cell coordinates and section-specific spatial scale factors, with communication probabilities constrained by physical distance (distance.use = TRUE, interaction range 250 µm, contact range 100 µm, scale.distance 0.05). To compare conditions, per-section results were aggregated and pathways were assessed for consistency across replicates. CellChat permutation p-values establish the existence of an interaction within a dataset rather than a significant between-condition difference, so condition differences were interpreted descriptively and required consistency across both nicotine versus both saline animals without overlap. Pathway-level information flow and sender-receiver structure were examined with rankNet, netVisual_aggregate, and netAnalysis functions. A parallel condition-level analysis combining biological replicates within each condition was performed for cross-comparison, and pooled-versus-per-section results were reconciled to guard against pooling artifacts.

### Software and data availability

Pseudobulk results for whole tissue and ROI comparisons are found in table S1. Raw and processed sequencing data have been deposited in GEO.

## Results

### Unsupervised clustering and spatial mapping of transcriptomic signatures in piriform cortex, isocortex and anterior olfactory nucleus

To examine how nicotine alters gene expression across the cellular and spatial organization of the anterior olfactory cortex including piriform cortex, male mice received a single intraperitoneal injection of nicotine (0.2 mg/kg) or saline (N=2 per condition), brains were collected 24 h later, and coronal sections spanning the anterior olfactory nucleus (AON), piriform cortex (PIR), and adjacent isocortex were profiled by Visium HD (Fig. 1A). Nucleus-based segmentation using H&E staining yielded 450,785 cells after quality control filtering across the four sections. Integration and clustering resolved 18 populations. To identify cell types in the AON, piriform cortex and surrounding isocortex, cells were annotated using the Allen Institute’s MapMyCells tool, which maps cells to the whole-mouse-brain taxonomy (23). Cell-type analysis identified cell types that occupied their expected anatomical positions including isocortical intratelencephalic (IT), extratelencephalic, corticothalamic excitatory neurons, olfactory excitatory neurons of the piriform/entorhinal (L2/3 IT PIR/ENT) and AON/taenia tecta (IT AON/TT) cortex, interneurons, and glial and vascular cells (Fig. 1B,C). Cell types were supported by canonical markers identified by FindAllMarkers (Fig. 1D), and *Ndst4* marked the ventral piriform domain (Fig. S1B).

**Figure 1.**
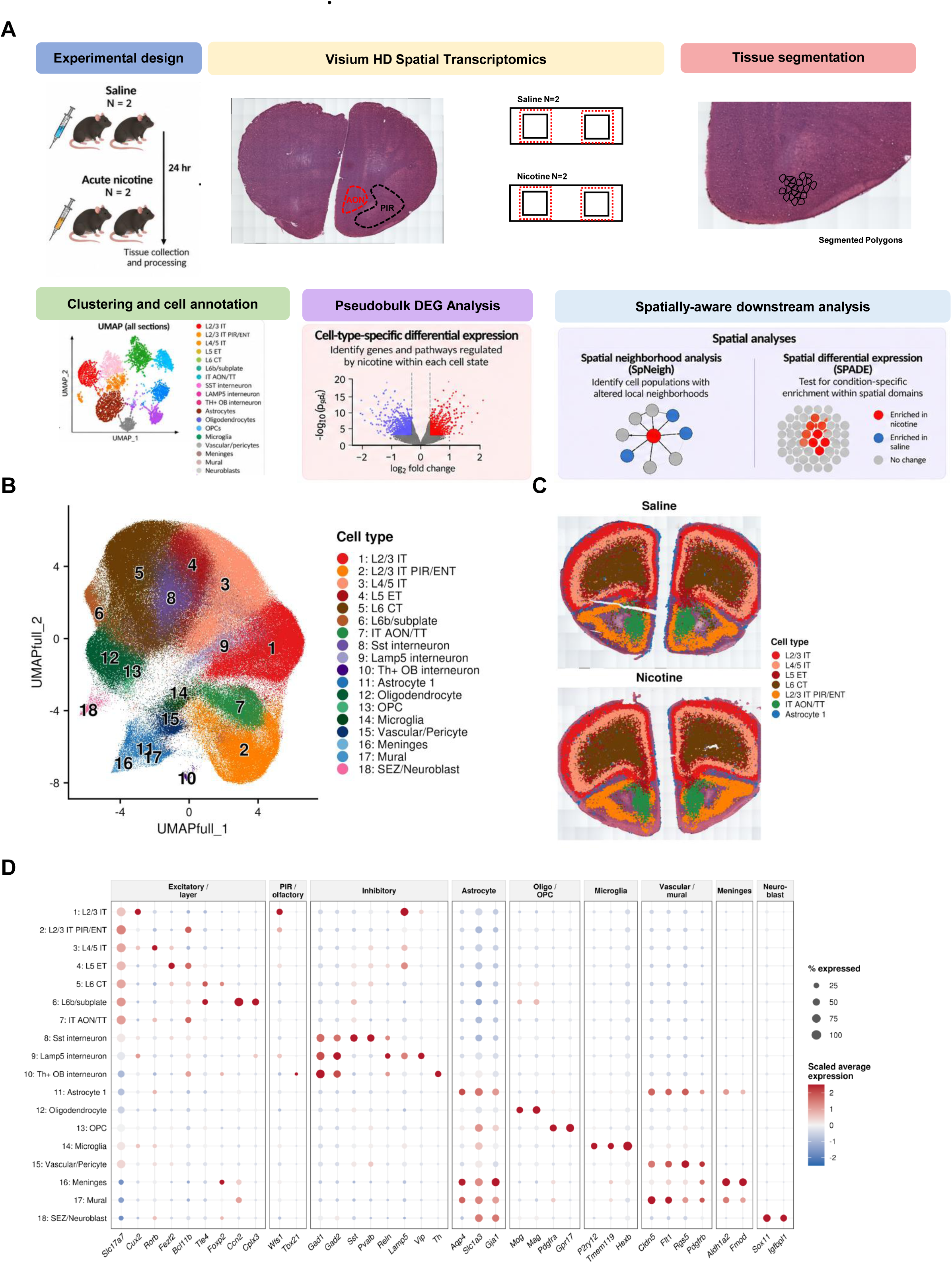
Spatial transcriptomic map of the anterior olfactory cortex and adjacent isocortex. **(A**) Experimental design. Adult male C57BL/6 mice received a single intraperitoneal injection of nicotine (0.2 mg/kg) or saline (N = 2 per condition). Brains were collected 24 h later, and one coronal section per animal spanning the anterior olfactory nucleus (AON), piriform cortex (PIR) and adjacent isocortex was profiled by Visium HD. Transcripts were assigned to individual cells by nucleus-based segmentation of the H&E image (Space Ranger v4.0.1). (**B**) Spatial map of annotated cell types in a representative section. (**C**) UMAP of the Harmony-integrated dataset (450,785 cells from four sections), colored by cell type. Annotation combined Allen Brain Cell atlas mapping (MapMyCells, 10x whole-mouse-brain taxonomy) with canonical marker genes. (**D**) Dot plot of marker genes for each cell type identified by FindAllMarkers. Dot size shows the fraction of cells expressing the gene; color shows scaled average expression. IT, intratelencephalic; ENT, entorhinal cortex; TT, taenia tecta.

### Acute nicotine selectively alters gene expression, and synapse signaling-associated gene signatures in olfactory cortical neurons

To test for differential expression within each cell type we used pseudobulk profiles summed per animal, treating animal rather than cell as the unit of replication (Squair et al., 2021). The response was concentrated in olfactory excitatory neurons (Fig. 2A). L2/3 IT PIR/ENT neurons had the most differentially expressed genes (DEGs; 78 total, 50 upregulated, 28 downregulated; FDR < 0.05), followed by IT AON/TT neurons (20). Isocortical IT neurons had fewer (14 in L4/5 IT, 13 in L2/3 IT), followed by *Sst* interneurons (7), and remaining populations had at most one gene. In piriform neurons, the largest changes included increased *Atp2b4*, encoding the plasma-membrane calcium ATPase PMCA4, together with *Abi3bp*, and *Marcksl1*, and decreased *Penk* (Fig. 2B). PMCA4 extrudes calcium from neurons and shapes the duration of activity-dependent calcium signals, *Abi3bp* encodes an extracellular-matrix protein and marks piriform pyramidal neurons (12), and *Marcksl1* encodes a protein kinase C substrate that regulates the actin cytoskeleton. Together, these changes suggest that 24 h after nicotine, piriform neurons had adjusted their calcium-handling and structural programs, while *Penk*, which encodes the opioid peptide precursor proenkephalin, was reduced.

**Figure 2.**
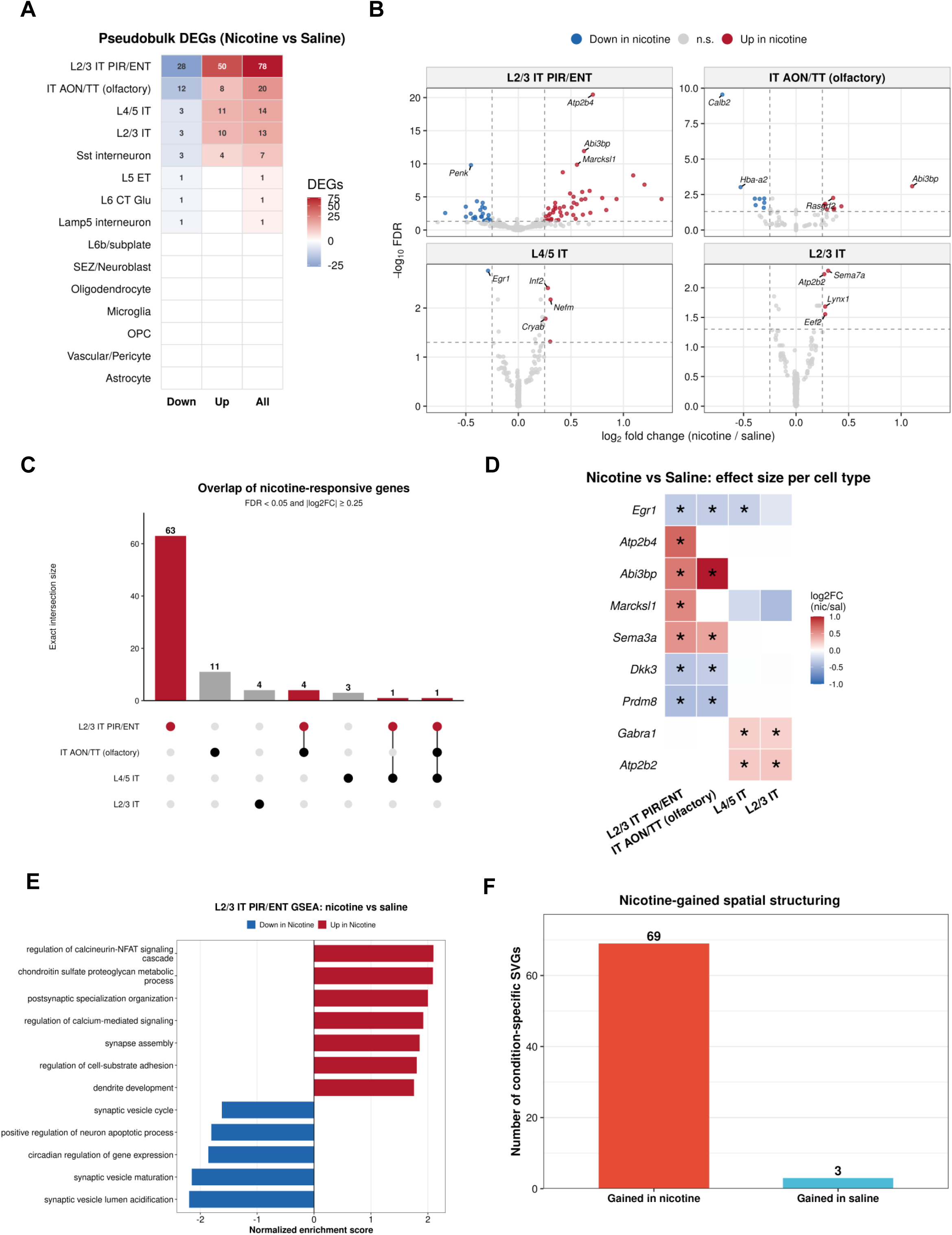
Nicotine’s transcriptional effects are concentrated in olfactory excitatory neurons. (**A**) Number of differentially expressed genes (DEGs; nicotine vs saline, FDR < 0.05) in each cell type. Counts come from pseudobulk DESeq2 analysis with animal as the unit of replication (one aggregated profile per animal per cell type; N = 2 per condition). (**B**) Volcano plot of nicotine-induced changes in L2/3 IT PIR/ENT neurons (apeglm-shrunken log2 fold changes). Selected genes, including *Atp2b4*, *Abi3bp*, *Marcksl1* and *Penk*, are labeled. (**C**) Overlap of DEGs across responsive cell types using UpSet plot (**D**) Log2 fold changes of DEGs shared by two or more populations, including *Egr1*, which was reduced in L2/3 IT PIR/ENT, IT AON/TT and L4/5 IT neurons. (**E**) Gene set enrichment analysis (gseGO, Biological Process) of whole-tissue L2/3 IT PIR/ENT neurons, with genes ranked by Wald statistic after excluding mitochondrial and ribosomal genes. Terms shown have adjusted p < 0.05; color shows direction (normalized enrichment score). (**F**) Genes spatially variable (SPARK-X, adjusted p < 0.05) in both nicotine sections and neither saline section (nicotine-gained, 69 genes), or the reverse (saline-gained, 3 genes).

The response was largely cell-type specific (Fig. 2C). *Egr1* was the only gene changed in both olfactory and isocortical neurons, and it was reduced in L2/3 IT PIR/ENT, IT AON/TT, and L4/5 IT neurons (Fig. 2D). Because nicotine-induced immediate-early gene expression normally returns to baseline within hours of a single injection (9), reduced *Egr1* at 24 h reflects a lasting change rather than the acute response. The two olfactory populations shared changes in *Sema3a*, *Dkk3*, *Prdm8*, and *Abi3bp*, whereas the two isocortical populations shared increases in *Gabra1*, a member of the KEGG nicotine addiction pathway, and *Atp2b2* (Fig. 2D). Gene set enrichment analysis (GSEA) of piriform neurons showed coordinated up-regulation of calcineurin-NFAT signaling, calcium-mediated signaling, chondroitin sulfate proteoglycan metabolism, postsynaptic specialization organization, synapse assembly, cell-substrate adhesion, and dendrite development, and down-regulation of synaptic vesicle cycling, maturation, and acidification, regulation of neuronal apoptosis, and circadian regulation of gene expression (Fig. 2E). Ligand-receptor inference (33) is presented descriptively in Supplementary Fig. S2A,B. These effects indicate that, 24 h after a single injection, nicotine’s transcriptional effects were concentrated in olfactory excitatory neurons and centered on calcium-handling, synapse-organizing, and extracellular-matrix programs, with reduced expression of the immediate-early gene *Egr1*.

Nicotine also altered spatial patterning. Sixty-nine genes were spatially variable in both nicotine sections and neither saline section, compared with three genes showing the converse pattern (Fig. 2F). These included *Klhl14*, which was also differentially expressed in piriform neurons and, with *Marcksl1*, was one of two genes whose pattern across cell-type domains differed between conditions (Fig. S1A). Together, these findings showed that nicotine’s effects were concentrated in, and spatially organized within, olfactory cortex, so we next examined a piriform/olfactory region of interest at higher cellular resolution.

### A Cartpt/Penk excitatory population carries the nicotine response in the piriform/olfactory region

To resolve the olfactory response at higher cellular resolution, we delineated a region of interest spanning PIR/ lateral entorhinal, and AON/TT cortex with a total of 158,478 cells (Fig. 3A,B). Reclustering separated excitatory neuronal states including a PIR-ENTl population co-expressing *Cartpt* and *Penk*, PIR-ENTl/AON glutamatergic neurons, AON-TT *Calb1/Calb2* and *Igfbp6/Crym* populations, L6b/endopiriform-claustrum glutamatergic neurons, and cortical IT neurons, together with interneurons and glia (Fig. 3B,C). The *Cartpt/Penk* population formed a continuous ribbon along piriform and lateral entorhinal cortex (Fig. 3A) and expressed markers of piriform superficial pyramidal cells (*Abi3bp, Slc30a3, Cux1*) while lacking semilunar-cell markers (*Reln, Rorb*, *Tcerg1l*; Fig. 3C) consistent with the *Cartpt*-expressing pyramidal supertype enriched in anterior piriform cortex (12).

**Figure 3.**
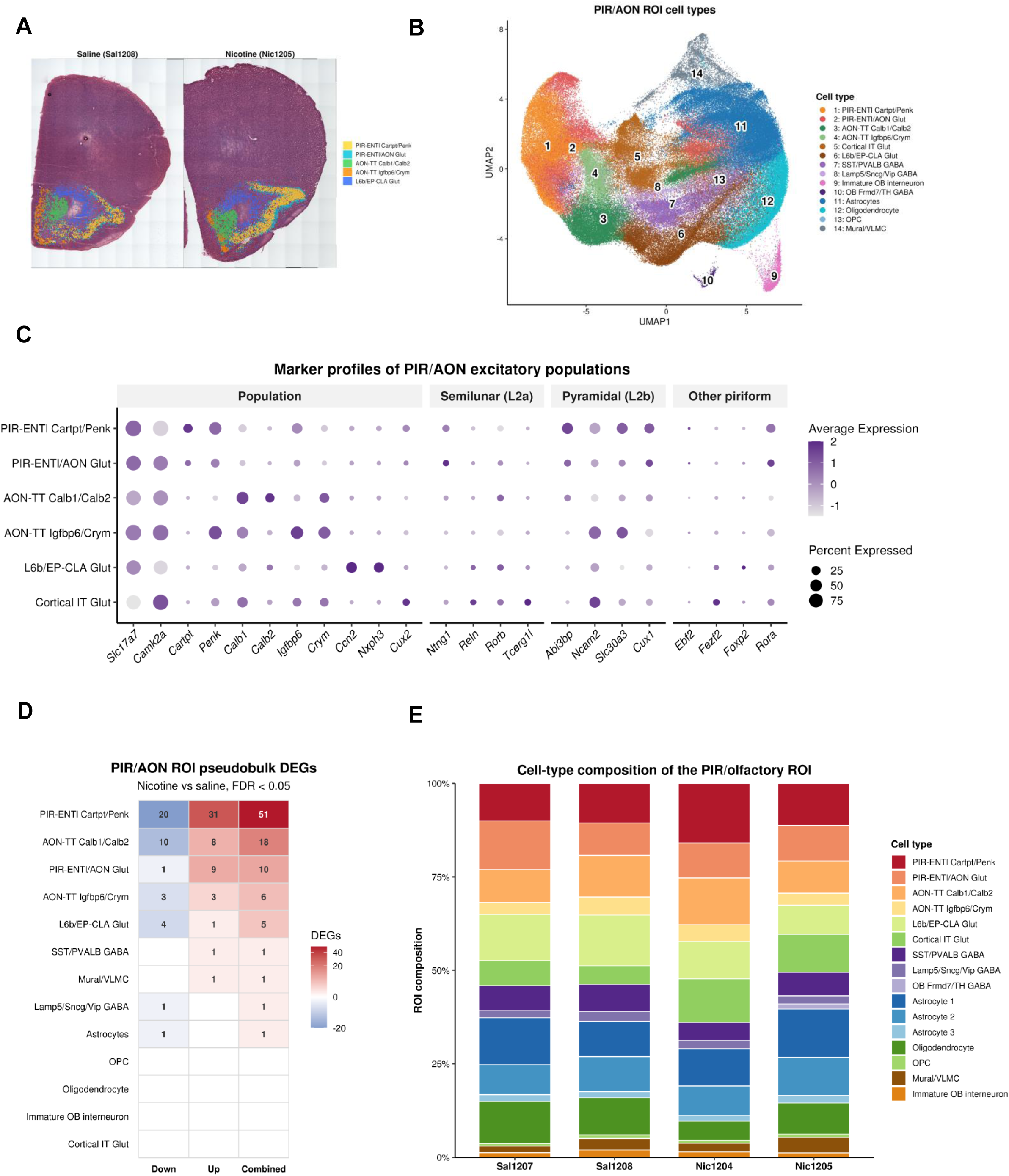
A *Cartpt/Penk* superficial pyramidal population carries the nicotine response in the piriform/olfactory region. (**A**) Spatial map of the region of interest (ROI) spanning PIR, lateral entorhinal cortex (ENTl) and AON/TT. *Cartpt/Penk* neurons form a continuous band along piriform and lateral entorhinal cortex for representative sections (**B**) UMAP of reclustered ROI cells (158,478 cells; Harmony integration, 30 principal components, resolution 0.4), colored by population. (**C**) Dot plot of markers distinguishing ROI populations. *Cartpt/Penk* neurons express superficial pyramidal cell markers (*Abi3bp*, *Slc30a3*, *Cux1*) and lack semilunar cell markers (*Reln*, *Rorb*, *Tcerg1l*). (**D**) Number of DEGs (FDR < 0.05) in each ROI population, from pseudobulk DESeq2 analysis with animal as the replicate (unshrunken estimates). (**E**) Cell-type proportions by condition. No population differed between nicotine and saline (propeller, all FDR > 0.05).

Within the region, PIR-ENTl *Cartpt/Penk* neurons had the most DEGs (51; 31 up, 20 down), followed by AON-TT Calb1/Calb2 (18) and PIR-ENTl/AON glutamatergic neurons (10); other populations had at most six, and astrocytes had one (Fig. 3D). Cell-type proportions did not differ between conditions (propeller, all FDR > 0.05, Fig. 3E), indicating that the transcriptional response was not driven by shifts in cell composition.

### Cartpt/Penk neurons up-regulate synaptic and extracellular-matrix programs

The *Cartpt/Penk* response (Fig. 4A) included increases in *Atp2b4, C1ql3, Ano3*, and *Adcyap1* and decreases *in Necab1, Egr1, Ppfia4, Bhlhe40, Tafa2*, and *Gpr88* (Fig. 4B). The defining neuropeptide genes themselves were not significantly changed (*Penk* log2 fold change −0.41; *Cartpt* log2 fold change −0.52, both FDR > 0.9) (Table S1). Enriched programs were up-regulation of chondroitin sulfate and proteoglycan metabolism, cell-substrate adhesion, neurotransmitter receptor regulation, postsynaptic specialization organization, synapse assembly, cell junction assembly, and dendrite development, and down-regulation of RNA splicing, mRNA processing, dendrite extension and cardiolipin and phosphatidylglycerol metabolism (Fig. 4C). The response thus overlapped the whole-tissue piriform response, including increased *Atp2b4*, decreased *Egr1*, and enrichment of synapse-assembly and proteoglycan programs, while localizing it to a spatially defined neuronal compartment. Consistent with the receptor-regulation signature, a test of the KEGG nicotine addiction gene set showed modest coordinated up-regulation in both whole-tissue piriform neurons and *Cartpt/Penk* neurons (Fig. 4C) (NES = 1.58, P = 0.02 in each), driven mainly by AMPA and NMDA receptor subunits (*Gria1-3, Grin2a, Grin2b*), GABA-A receptor subunits, and *Chrna4* (Fig. 4D).

**Figure 4.**
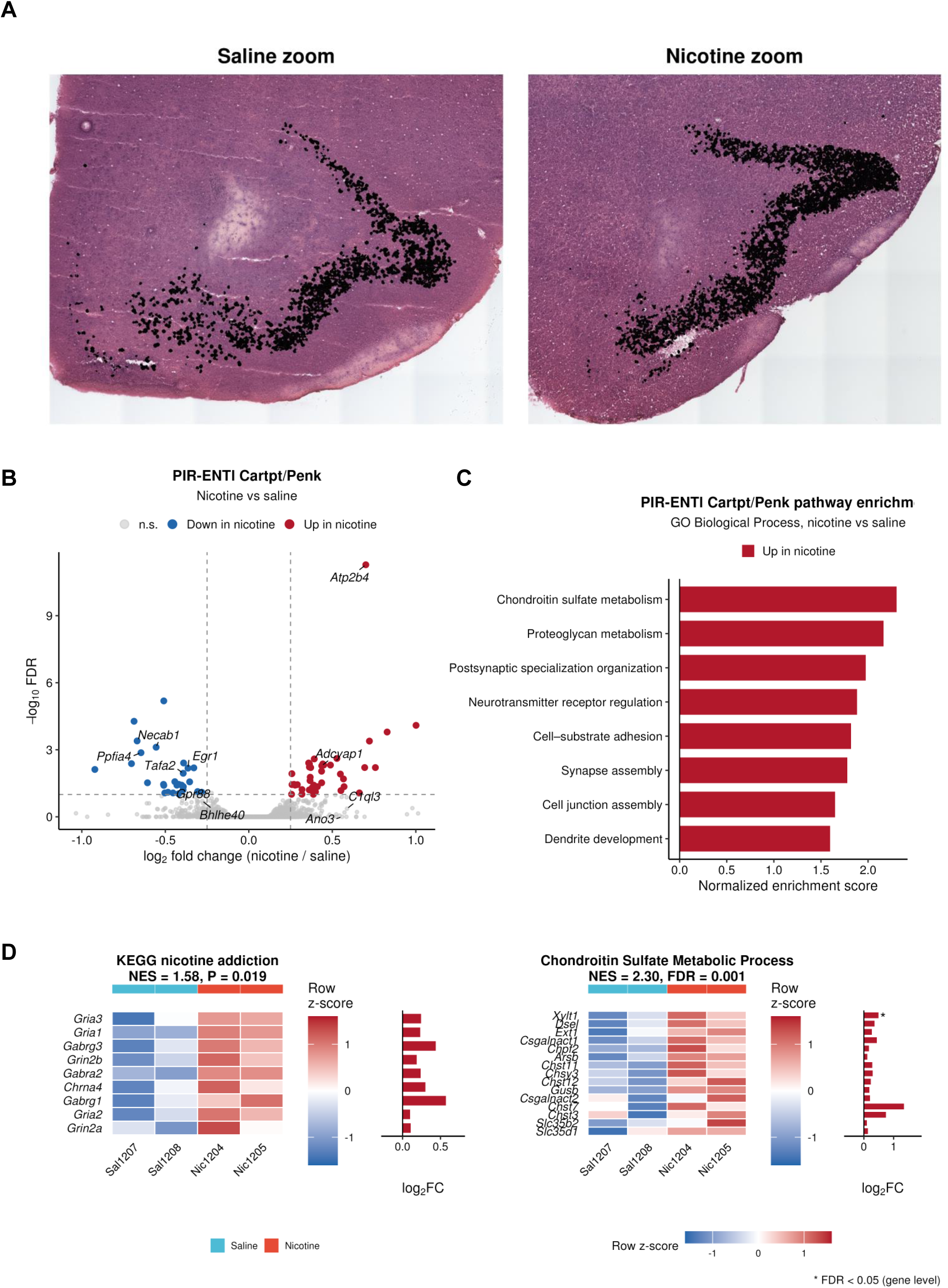
*Cartpt/Penk* neurons upregulate synaptic, receptor and extracellular-matrix programs after nicotine. (**A**) Volcano plot of nicotine-induced changes in *Cartpt/Penk* neurons: 51 DEGs, 31 up and 20 down (FDR < 0.05). (**B**) Log2 fold changes of top DEGs, including increased *Atp2b4*, *C1ql3*, *Ano3* and *Adcyap1*, and decreased *Necab1*, *Egr1*, *Ppfia4*, *Bhlhe40*, *Tafa2* and *Gpr88*. (**C**) Gene set enrichment analysis (gseGO, Biological Process) for *Cartpt/Penk* neurons (adjusted p < 0.05). (**D**) Left: enrichment of the KEGG nicotine addiction gene set (mouse orthologs of hsa05033; fgsea) in whole-tissue piriform neurons and *Cartpt/Penk* neurons (NES = 1.58, p = 0.02 in each). Leading-edge genes of the KEGG nicotine addiction set, Right: Leading-edge genes for Chondroitin Sulfate Metabolic Process

Ligand-receptor inference identified glutamate as the dominant input to *Cartpt/Penk* neurons, per-animal communication probably for the strongest glutamate and PTN ligand-receptor pairs targeting *Cartpt/Penk* neurons, which some pairs differing between saline and nicotine animals(S3B). Neither channel differed consistently between nicotine and saline animals, so these communication results are presented descriptively. Their presence, together with the extracellular-matrix and synapse-assembly programs of *Cartpt/Penk* neurons, nonetheless pointed to the surrounding astrocytic environment, which we examined next.

### Nicotine lowers *Sparcl1* in the astrocyte-bordered tissue adjacent to Cartpt/Penk neurons

Astrocytes lined the *Cartpt/Penk* ribbon and were interspersed with flanking PIR-ENTl/AON glutamatergic neurons (Fig. 5A). Because the *Cartpt/Penk* program was enriched for synapse-assembly and extracellular-matrix terms, processes strongly shaped by astrocyte-secreted proteins, we asked whether expression in the surrounding tissue varied with distance from *Cartpt/Penk* neurons and whether nicotine changed that relationship. We binned PIR-ENTl/AON glutamatergic cells by distance to the nearest *Cartpt/Penk* cell and, for each gene, tested whether the shape of the expression-distance curve differed between conditions (Fig. 5B). The strongest genome-wide effect was for *Sparcl1* (Fig. 5D), which encodes the synaptogenic matricellular protein hevin (condition x distance FDR = 0.011); *Pcp4*, *Btbd17*, and *Rora* also passed FDR < 0.05.

**Figure 5.**
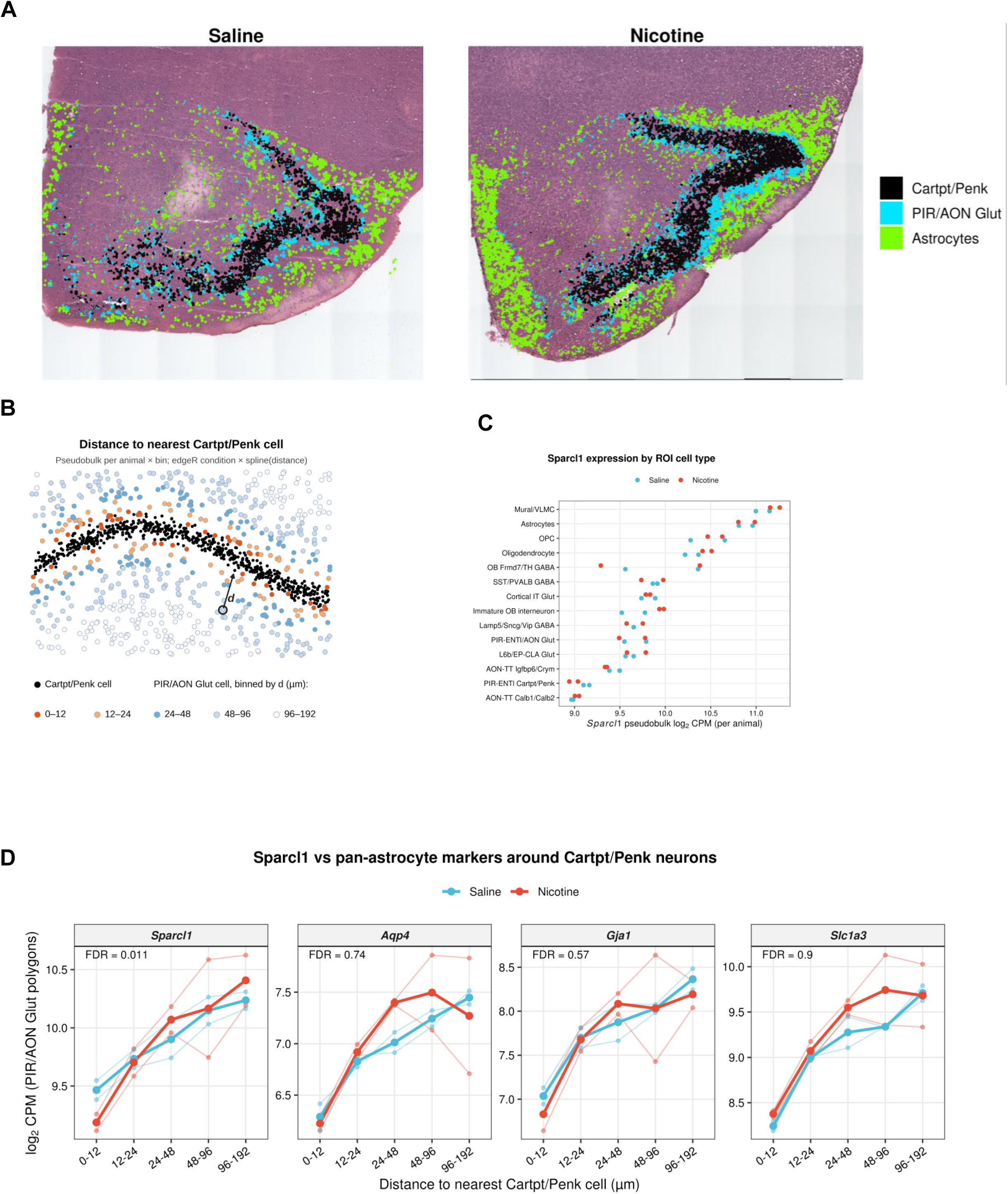
Nicotine lowers *Sparcl1* in the tissue immediately bordering *Cartpt/Penk* neurons. **(A)** Spatial distribution of astrocytes relative to the *Cartpt/Penk* band and the flanking PIR-ENTl/AON glutamatergic neurons in a representative section. (**B**) Analysis schematic. Each PIR-ENTl/AON glutamatergic cell was assigned to a bin by its distance to the nearest *Cartpt/Penk* cell: 0–12, 12–24, 24–48, 48–96 and 96–192 µm). Counts were aggregated per animal and bin, and fit in edgeR with natural cubic splines of log2 distance (df = 2). A quasi-likelihood F-test of the condition × spline terms tested whether the expression–distance relationship differed between nicotine and saline. (**C**) *Sparcl1* expression across ROI cell types. Expression is highest in mural/vascular-leptomeningeal cells and astrocytes and lowest in *Cartpt/Penk* neurons. (**D**) Genome-wide results of the condition × distance test (10,861 genes). *Sparcl1* is the top-ranked gene (FDR = 0.011); *Pcp4*, *Btbd17* and *Rora* also pass FDR < 0.05. (E) Expression versus distance from the *Cartpt/Penk* band for *Sparcl1* and the pan-astrocytic markers *Aqp4*, *Gja1* and *Slc1a3*, by condition. In both conditions *Sparcl1* rises with distance, and nicotine lowers it in the 0–12 µm bin. The astrocytic markers follow the same gradient with no nicotine interaction (FDR = 0.74, 0.57 and 0.90). Points are animal × bin pseudobulk values; lines are model fits.

*Sparcl1* was glia-enriched in this tissue, highest in mural/vascular leptomeningeal cells and astrocytes and lowest in *Cartpt/Penk* neurons (Fig. 5C). In both conditions it rose with distance from the *Cartpt/Penk* ribbon, away from the border of the pyramidal layer. Nicotine lowered it further in the bin nearest the ribbon (0-12 µm), and the curves converged by 24-48 µm (Fig. 5E). The pan-astrocyte markers *Aqp4, Gja1*, and *Slc1a3* followed the same distance gradient in both conditions but showed no nicotine interaction (FDR = 0.74, 0.57, and 0.90 respectively; Fig. 5E). The baseline gradient therefore reflects the distribution of astrocytic material around the dense *Cartpt/Penk* layer, whereas the nicotine effect is not explained by a general loss of astrocytic content.

The *Sparcl1* gradient itself was present within individual sections by an independent boundary-based analysis (31) (boundary-proximity trend FDR < 8.1 x 10^-4 in each section) (supplementary files to come). The effect was strongest in glutamatergic polygons that also contained astrocytic transcripts (P = 1.2 x 10^-5) and weaker but detectable in polygons without detectable astrocyte markers (P = 0.011), consistent with contributions from astrocytic *Sparcl1*. The distance interaction was significant in edgeR. By contrast, astrocyte transcriptomes themselves were largely unchanged. Together, these results indicate that acute nicotine reduces a glia-enriched synaptogenic signal specifically in the tissue immediately adjacent to the responding *Cartpt/Penk* neurons.

## Discussion

Using single-cell resolution spatial transcriptomics, we characterized the transcriptional response to acute nicotine across the anterior olfactory cortex and adjacent isocortex. We found that the response was neuronal and regionally selective, concentrated in olfactory excitatory neurons. The strongest response localized to a spatially defined population of piriform/entorhinal neurons marked by the reward-associated neuropeptide genes *Cartpt* and *Penk*, which up-regulated calcium-handling, synapse assembly, and extracellular-matrix programs. Additionally, nicotine lowered *Sparcl1*, which encodes the astrocyte-secreted synaptogenic protein hevin, specifically in the tissue immediately adjacent to these neurons, without a corresponding change in pan-astrocytic markers. Together, these findings indicate that a single nicotine exposure produces a lasting, spatially organized transcriptional response centered on one olfactory-cortical neuronal population and its immediate surroundings.

### Nicotine engages a cholinergic-regulated olfactory learning circuit

The concentration of the response in olfactory cortex fits the strong cholinergic control of this region, which supports olfactory learning and memory (5,6,34). Blocking nicotinic receptors in the olfactory bulb impairs discrimination of perceptually similar odors (5), and systemic nicotine induces *c-fos* mRNA in piriform cortex (9). Nicotine’s actions on this system have been studied mainly at the level of behavior, where chronic oral nicotine improves olfactory memory and reduces neuronal apoptosis in piriform cortex in a TRPA1-dependent manner (10). Notably, regulation of neuronal apoptosis was among the programs reduced in piriform neurons in our data. Our data extend these observations to cell-type and spatial resolution. In our previous study, bulk RNA sequencing of the hippocampus after the same exposure identified differentially expressed genes associated with nicotine-related smoking initiation GWAS loci (19). The present approach extends that work by resolving which cell types and tissue locations carry the response, in a different forebrain region. Together, these observations suggest that nicotine preferentially engages forebrain circuits specialized for associative learning, including the olfactory cortex.

Nicotine is also a potent modulator of glutamatergic signaling, increasing glutamate release through presynaptic nicotinic receptors (13) and facilitating glutamatergic potentiation (14), and the *Cartptp/Penk* response included up-regulation of glutamate receptor genes. Complementary *in vivo* imaging also implicates piriform cortex in nicotine’s effects on glutamatergic networks. Using glutamate-weighted chemical exchange saturation transfer (gluCEST) MRI, (11) found that seven days of nicotine (10 mg/kg/day, i.p.) broadly attenuated glutamate-associated connectivity in the mouse brain and disrupted a sensory-limbic module linking the lateral geniculate nucleus, amygdala, and piriform cortex. That study differs from ours in dose, duration, and measurement, reporting network-level glutamate-weighted contrast rather than cell-type-resolved transcription. Nonetheless, a network-level glutamatergic signal in piriform cortex after repeated nicotine and the glutamate-receptor and synaptic programs we observed in piriform pyramidal neurons after a single dose suggest that glutamatergic signaling in piriform cortex may be a shared target of nicotine across timescales and methods.

### Nicotine’s response localizes to Cartpt/Penk superficial pyramidal neurons

*Cartpt/Penk* neurons were identified as the most responsive population, a subgroup that has a defined place in piriform circuity. The *Cartpt/Penk* neuronal population expressed markers consistent with layer 2b superficial pyramidal cells and lacked semilunar-cell markers, matching a *Cartpt*-expressing pyramidal supertype found enriched in the anterior piriform cortex (12). Superficial pyramidal cells differ from semilunar cells in two ways that bear on nicotine’s effects. They are recurrently connected, forming the associational network thought to store odor representations, whereas semilunar cells relay bulbar input to them without recurrent connections (8). These neurons also project beyond the piriform cortex, including to the orbitofrontal and medial prefrontal cortex (7). Disconnecting piriform from orbitofrontal cortex reduces relapse to fentanyl seeking in rats (35). Although this was not tested for nicotine, odors act as salient smoking cues, and cigarette odor increases craving in nicotine-dependent smokers (36). Our mice received nicotine by injection, without accompanying odor, so the relevance is not that nicotine is smelled but that the drug acts on the circuit that encodes odor associations; in smokers, nicotine and smoking-related odors occur together, so a drug acting on this circuit could influence how those odors are encoded. Nicotine’s strongest cortical response therefore falls on neurons within the circuitry that stores odor associations and sends them to frontal regions, and whether this affects odor memory or cue-related behavior remains to be tested.

These neurons are also defined by two neuropeptide genes with established roles in drug reward. CART was named for its induction in the striatum by cocaine and amphetamine (37). CART is expressed prominently in mesocorticolimbic targets including piriform cortex (38,39). Additionally, nicotine regulates CART in the mesocorticolimbic system with repeated dosing (40). Proenkephalin-derived peptides are required for nicotine reward in mice (41). In our study, however, neither *Cartpt* nor *Penk* changed significantly in these neurons after a single dose, their synaptic and receptor programs changed instead. The neuropeptide identity of this population therefore marks a cell type rather than indicating engagement of reward signaling, and whether these neurons contribute to nicotine-related behavior remains to be tested. Together, these features suggest that at 24 h, nicotine’s effect on this population is primarily synaptic and structural rather than relating to neuropeptides. Several features of the *Cartpt/Penk* program parallel prior nicotine studies. Nicotine self-administration remodels perineuronal nets, which are built from chondroitin sulfate proteoglycans, in the ventral tegmental area and orbitofrontal cortex (42), and down-regulation of cardiolipin and phosphatidylglycerol metabolism, lipids of the inner mitochondrial membrane, echoing nicotine-induced mitochondrial transcriptional changes in ventral tegmental area neurons (21). These studies support our gene set enrichment results which found up-regulation of chondroitin sulfate and proteoglycan metabolism. Coordinated up-regulation of glutamate receptor subunits likewise fits nicotine’s established facilitation of glutamatergic transmission (13).

### A local change in Sparcl1 at the border of responding neurons

Nicotine lowered *Sparcl1* only within about one cell diameter of the *Cartpt/Penk* neurons, whereas pan-astrocytic markers showed the same distance gradient in both conditions without a nicotine effect. This local effect is compatible with our whole-tissue spatial analyses, which addresses different questions. *Sparcl1* is spatially patterned in every section, so it cannot appear as a nicotine-gained spatially variable gene, and testing for differential spatial patterns across whole cell-type domains averages over the narrow band in which the change occurs. Injected nicotine reaches the entire brain section, so a drug acting directly on astrocytes would be expected to change *Sparcl1* throughout the tissue. Restriction of the change to the border of the responding neurons instead suggests that it depends on those neurons, for example through signals they release, although our data cannot identify the mechanism.

*Sparcl1* encodes hevin, an astrocyte-secreted protein that promotes excitatory synapse formation, in part by bridging neurexin-1α and neuroligin-1B (43). Its synaptogenic activity is antagonized by SPARC (44), and it also acts independently of neurexins and neuroligins (45). While *Cartpt/Penk* neurons up-regulated synapse-assembly and glutamate-receptor programs, a pro-synaptogenic signal at their border was reduced. One possibility is that lower hevin restrains the addition of excitatory synapses onto neurons that have just responded strongly, because *Sparc* expression did not change with distance or condition, the balance between hevin and its antagonist would shift away from synaptogenesis near the ribbon. Because the same neurons also up-regulated chondroitin sulfate and proteoglycan programs, the reduction may instead form part of a coordinated remodeling of the extracellular environment at this border. Hevin has recently been linked to drug-induced plasticity: in the nucleus accumbens, astrocytic hevin was required for cocaine place preference and cocaine-induced synaptic plasticity (46). Our observation extends the association between hevin and drug exposure in the olfactory cortex, but at the transcript level and at a single time point. Hevin levels also matter for memory: raising astrocytic hevin in the hippocampus improved memory in aged and Alzheimer’s-model mice by altering synaptic composition (47), indicated that hevin supplied by astrocytes can shape synapses relevant to cognition. Whether a local reduction at the border of piriform pyramidal neurons has functional consequences remains to be tested. These findings suggest that nicotine alters the local synaptogenic environment of the neurons it most strongly affects, a hypothesis that in situ validation can directly test (we are working on it-lambo)

The cellular source of the change is unresolved. Astrocytes express functional nicotine receptors (17), but the effect was also present, more weakly, in glutamatergic polygons lacking detectable astrocyte transcripts. Neurons can express *Sparcl1*, and nucleus-based segmentation assigns some transcripts to neighboring cells. We therefore describe the change as occurring in the tissue bordering *Cartpt/Penk* neurons rather than in a specific cell type. These results can be compared with our earlier finding that the same dose increased astrocyte morphological complexity 24 h after injection, with AKT2 modulating this response (19). Here, astrocyte transcriptomes in olfactory cortex were largely unchanged, yet *Sparcl1* changed locally at the border of responding neurons. As Visium HD does not capture astrocyte morphology and the two studies examined different regions, we cannot determine whether the local *Sparcl1* change accompanies the process remodeling we observed previously.

### Limitations and future directions

Since this study measured transcripts in two animals per condition and did not assess behavior, we are able to generate hypotheses meant to be tested rather than conclusions. First, the synaptic, receptor, and extracellular-matrix programs of *Cartpt/Penk* neurons 24 h after a single dose may reflect a lasting adaptation of piriform associative circuitry, predicting corresponding changes in synaptic proteins or excitatory synapse density near the *Cartpt/Penk* layer. Second, the local *Sparcl1* reduction may depend on the responding neurons, tracking the time course of the neuronal response and is absent where *Cartpt/Penk* neurons are not engaged. Time-course and repeated-exposure designs would directly test this. Third, because piriform cortex supports odor learning and odors are salient smoking cues, these changes could influence how odor cues become associated with nicotine, which could be tested with future behavioral experiments. Only male mice, a single dose, and a single post-injection interval (24 h) were studied. Our earlier bulk and morphological analyses focused on a different region, so direct comparison between the studies is limited.

## Conclusion

A single nicotine exposure produced a lasting, neuron-centered transcriptional response in olfactory cortex that localized to *Cartpt/Penk* superficial pyramidal neurons and was accompanied by a local reduction of *Sparcl1* at their border. These transcriptional changes occurred without detectable treatment-associated shifts in overall cell-type composition, suggesting that acute nicotine primarily alters cellular state and local tissue organization rather than gross cellular abundance. The spatial restriction of these effects indicates that even a single nicotine exposure can leave a selective molecular imprint on associative sensory circuitry 24 h later, rather than producing a uniform response across the forebrain. These results identify specific cell types, genes, and tissue locations for targeted validation and illustrate how spatial transcriptomics can reveal drug responses organized by tissue position.

---

**Supplementary Figure S1.**
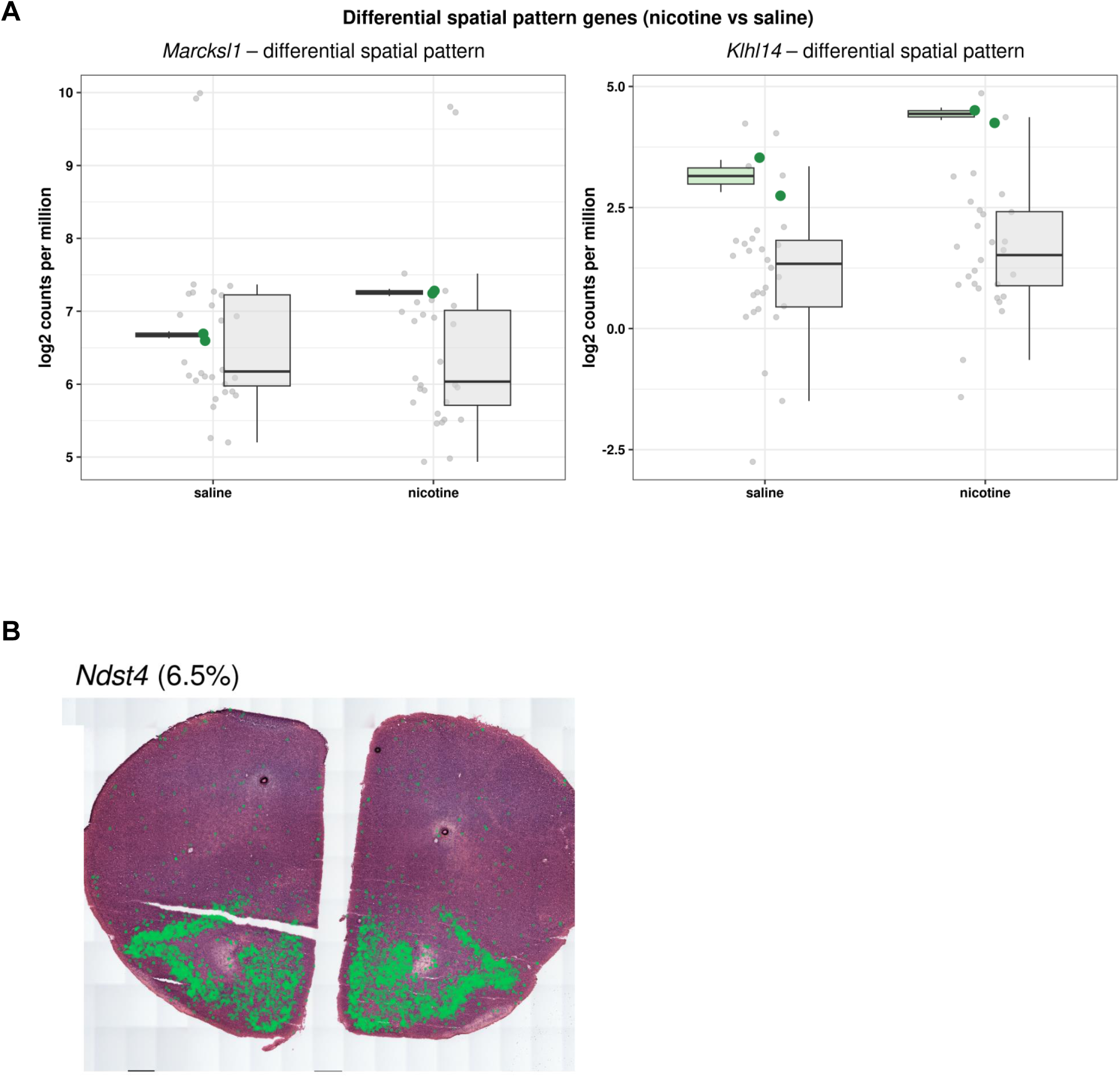
Spatial markers and differential spatial patterns. **(A)** Genes whose expression pattern across cell-type domains differs between conditions (DESpace framework; edgeR quasi-likelihood test of domain × condition interaction, FDR < 0.05): *Klhl14* and *Marcksl1*. (**B**) Spatial expression of *Ndst4*, which marks the ventral piriform domain.

**Supplementary Figure 2.**
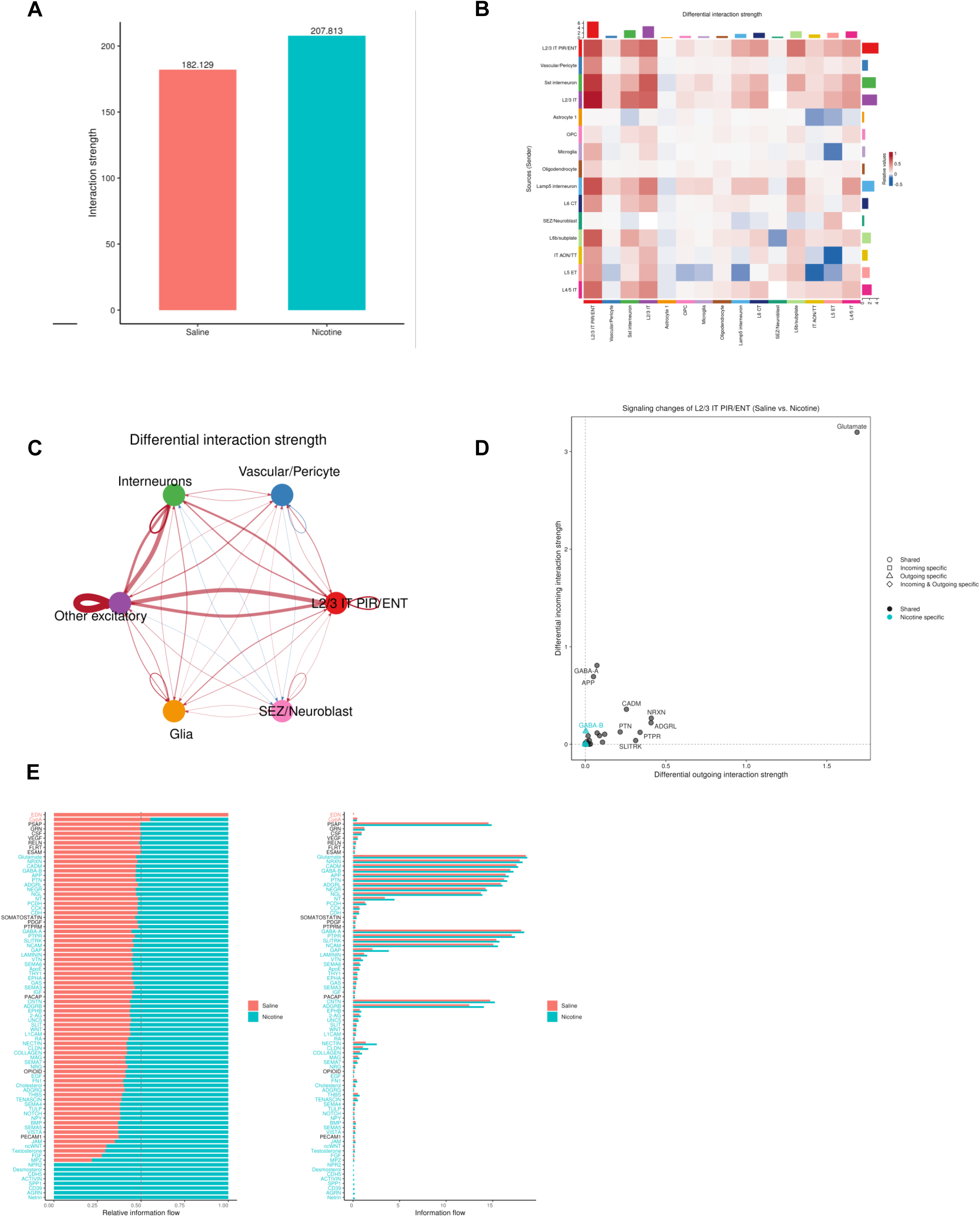
Inferred cell–cell communication in saline-and nicotine-treated olfactory cortex. (**A**) Total inferred interaction strength across all cell types in saline (182.1) and nicotine (207.6) conditions. (**B**) Heatmap of differential interaction strength (nicotine minus saline) between sender (rows) and receiver (columns) cell types. Red indicates stronger communication in nicotine, and blue indicates stronger communication in saline. Bars on the top and right show the summed change in incoming and outgoing signaling for each cell type. (**C**) Differential interaction strength among aggregated cell groups (L2/3 IT PIR/ENT, other excitatory neurons, interneurons, glia, vascular/pericyte and SEZ/neuroblast). Edge width is proportional to the magnitude of the change. Red edges indicate increased communication in nicotine, and blue edges indicate decreased communication. (**D**) Signaling changes in L2/3 IT PIR/ENT neurons. Each point is a signaling pathway, plotted by its differential outgoing (x-axis) and incoming (y-axis) interaction strength in nicotine relative to saline. Shape indicates whether the change is shared or specific to incoming or outgoing signaling. Color distinguishes pathways active in both conditions (black) from pathways detected only in nicotine (cyan). Glutamate signaling shows the largest increase in incoming interaction strength. (**E**) Information flow for each signaling pathway, summed across all cell pairs, in saline (red) and nicotine (cyan). Left: relative information flow. Right: absolute information flow. Pathway names are colored by the condition in which they are enriched

**Supplementary Figure 3.**
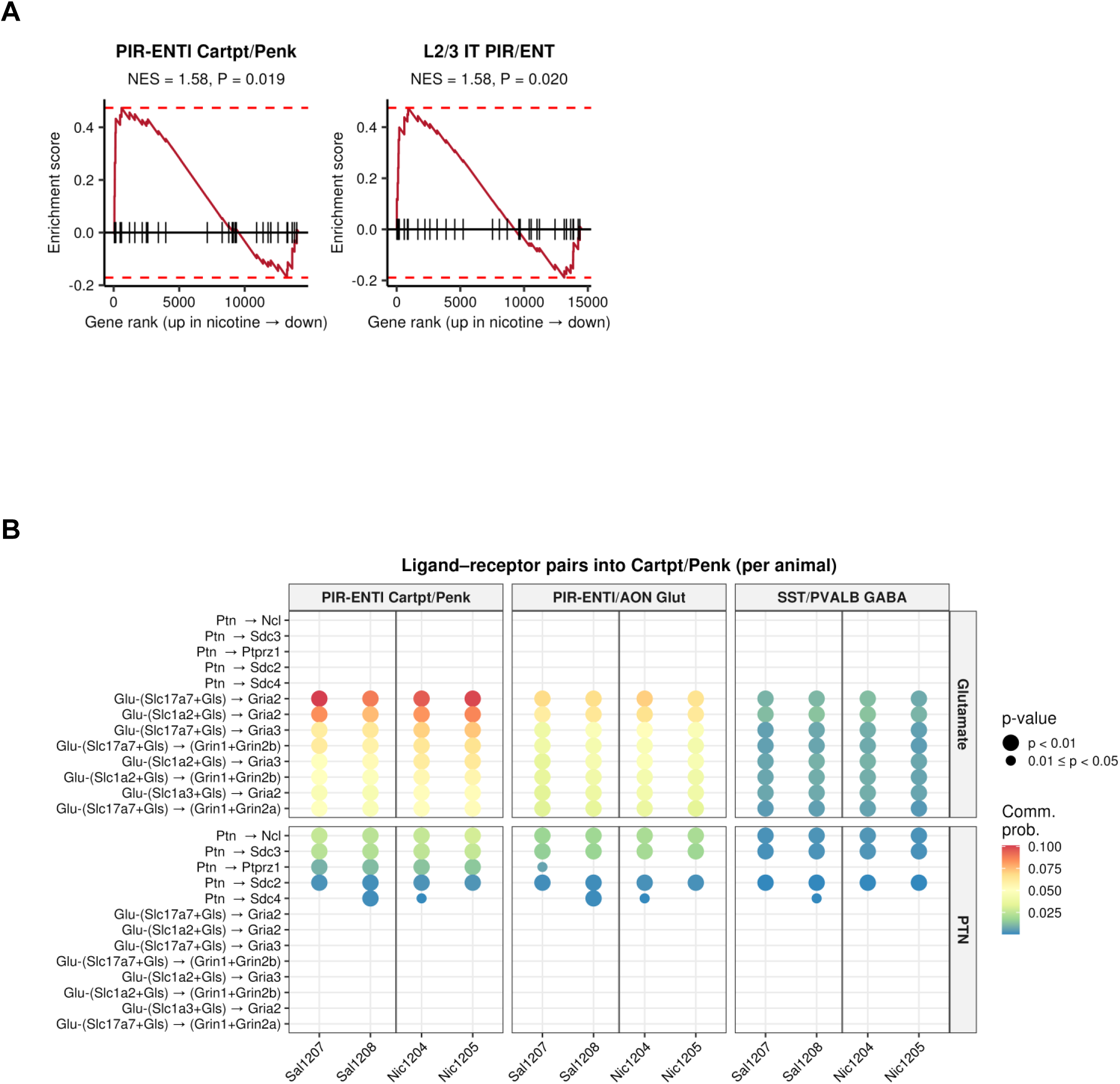
Nicotine addiction gene set enrichment and per-animal ligand–receptor inputs to *Cartpt/Penk* neurons. **(A)** Gene set enrichment plots for the KEGG nicotine addiction pathway (mouse orthologs of hsa05033). Genes are ranked by the DESeq2 Wald statistic for nicotine versus saline, from most up in nicotine (left) to most down (right). Results are shown for PIR-ENTl *Cartpt/Penk* neurons (left; NES = 1.58, P = 0.019) and whole-tissue L2/3 IT PIR/ENT neurons (right; NES = 1.58, P = 0.020). Vertical ticks mark the positions of pathway genes in the ranked list. The red curve is the running enrichment score, and dashed lines mark its maximum and minimum. Enrichment was tested with fgsea after excluding mitochondrial and ribosomal genes. (**B**) Communication probabilities for the strongest glutamate (top) and pleiotrophin (PTN, bottom) ligand–receptor pairs targeting *Cartpt/Penk* neurons, inferred separately for each animal with spatially constrained CellChat. For comparison, the same pairs are shown for two other receiving populations, PIR-ENTl/AON glutamatergic neurons and SST/PVALB GABAergic interneurons. Columns are individual animals (Sal1207 and Sal1208, saline; Nic1204 and Nic1205, nicotine). Dot color shows communication probability, and dot size shows the permutation p-value (large, p < 0.01; small, 0.01 ≤ p < 0.05). Pairs without a dot were not detected at p < 0.05. CellChat represents glutamate ligands as the combination of a vesicular transporter (*Slc17a6*/*Slc17a7*) and glutaminase (*Gls*).

